# Recruitment status and host-fish-limitation threats to endangered freshwater pearl mussel (*Margaritifera laevis*) in eastern Hokkaido, northern Japan

**DOI:** 10.1101/2023.05.19.541543

**Authors:** Kazuki Miura, Nobuo Ishiyama, Junjiro N. Negishi, Keita Kawajiri, Hokuto Izumi, Daisetsu Ito, Futoshi Nakamura

## Abstract

Recruitment failure is a major threat to freshwater mussel (Order Unionoida) populations worldwide. Assessments of the recruitment status and determining the bottleneck factors of mussel recruitment are crucial for preventing future declines in mussel populations. In this study, we investigated the recruitment status (i.e., size structure and the proportion of juveniles within a population) of the endangered freshwater pearl mussel *Margaritifera laevis* in 22 rivers in eastern Hokkaido, northern Japan. We also quantified the density of the host fish *Oncorhynchus masou masou* and examined the relationship between the proportion of juveniles and host-fish density to assess host-limitation threats to *M. laevis* recruitment. Our assessments showed that 13 (59.1%) rivers had no signs of recent recruitment within 10 years, with a low mean proportion of juveniles (mean:0.02 [range:0.00–0.09] fraction), indicating that these populations are threatened by sustained recruitment failure. The proportion of juveniles was positively associated with host-fish density, suggesting that host-fish limitation could be a bottleneck factor for *M. laevis* recruitment. These results highlight the urgent need for prompt conservation measures, including the enhancement of host-fish availability, to sustain *M. laevis* populations in the study region.

## Introduction

Freshwater mussels (Order Unionoida), with more than 800 described species, are widely distributed in various freshwater ecosystems worldwide, except Antarctica (Graf and Cummings 2007). They play key ecological roles in freshwater habitats, including modification of physical habitat characteristics, nutrient cycling alteration, and water purification (Gutiérrez et al. 2003; Vaughn and Hoellein 2018). However, they are among the most imperiled faunal groups of organisms (Lopes-Lima et al. 2018). The conservation of freshwater mussels is important for maintaining the structure and function of freshwater ecosystems (Miura et al. 2021b).

A recent review has highlighted that recruitment failure is a major threat to freshwater mussels (Ferreira-Rodríguez et al. 2019). Many populations have low or no recent recruitment, and the lack of young individuals is not necessarily associated with the population size (Österling et al. 2008; Geist 2010). If recruitment is not recovered, these populations will disappear and/or shrink with a long time lag because of their longevity (typically exceeding decades) (Haag and Rypel 2011; Hylander and Ehrlén 2013). Therefore, assessments of recruitment status, such as the size structure and proportion of young individuals within a population, are critical for evaluating actual population health and prioritizing the population targeted for conservation (Geist 2010; Sousa et al. 2015).

In addition to recruitment status assessments, determining the cause of recruitment failure is crucial for preventing future local extinction and/or population shrinkage of freshwater mussels (Österling et al. 2010; Strayer and Malcom 2012). The recruitment of freshwater mussels strongly depends on the availability of the host fish because they have a complex life cycle that necessarily parasitizes suitable host fish during their larval phase (Modesto et al. 2018). Thus, examining the influence of host-fish limitation using quantitative fish data is a vital first step in identifying the cause of recruitment failure in freshwater mussel species (Arvidsson et al. 2012; Stoeckl et al. 2015). However, such empirical studies have been limited to only a few mussel species and regions.

Freshwater pearl mussel, *Margaritifera laevis* (Haas, 1910), is distributed in streams in Sakhalin and Kamchatka, Russia and the Honshu and Hokkaido Islands of Japan (Sakai et al. 2022). This species has more than 80 years of longevity by region, and Masu salmon, *Oncorhynchus masou masou*, is a natural host (Kawajiri et al. 2021). The reproductive season for this species is mid-June to August (Akiyama 2007; Terui et al. 2015). *Margaritifera laevis* has experienced a significant population decline in Japan in recent decades and is listed as “Endangered” on the Red List of Japan (Ministry of the Environment of Japan 2020). Evaluation of the population status of this species is required to develop prompt planning for conservation measures (Miura et al. 2019a). Among the *M. laevis* distribution ranges in Japan, many *M. laevis* populations have been confirmed in the eastern part of Hokkaido (Miura et al. 2019b); however, recruitment status and its limiting factors have not been intensively evaluated.

Here, we aimed to clarify the recruitment status of *M. laevis* in eastern Hokkaido and the potential influence of host-fish limitations on recruitment failure. First, we determined the size structure and the proportion of young individuals within an *M. laevis* population in the 22 studied rivers. Second, we quantified the abundance of host fish in the streams and tested the relationship between mussel recruitment and host-fish availability.

## Materials and Methods

### Study site

The study was conducted in 22 rivers (sites) across eight watersheds, where several *M. laevis* populations were previously determined (Miura et al. 2019a, b), comprising >1500 km^2^ of eastern Hokkaido (Fig. 1). Thirteen (sites *a* to *m*) and nine (sites *n* to *v*) sites drain into the Nemuro Strait and the Pacific Ocean, respectively. No large dams or weirs inhibited the migration of host fish. The sites were not set up of each site upstream to ensure site independence. *Margaritifera kurilensis* (Zatravkin & Starobogatov, 1984) also occurred at study sites (Miura et al., 2019b). *Margaritifera kurilensis* has a different reproductive season (March to May) and suitable host-fish species (white-spotted char *Salvelinus leucomaenis leucomaenis* (Pallas, 1814) and perhaps Dolly Varden char *S. malma krascheninnikovi* Taranetz, 1933) with *M. laevis* (Kobayashi and Kondo 2009; Miura et al. 2021a).

**Fig. 1.**
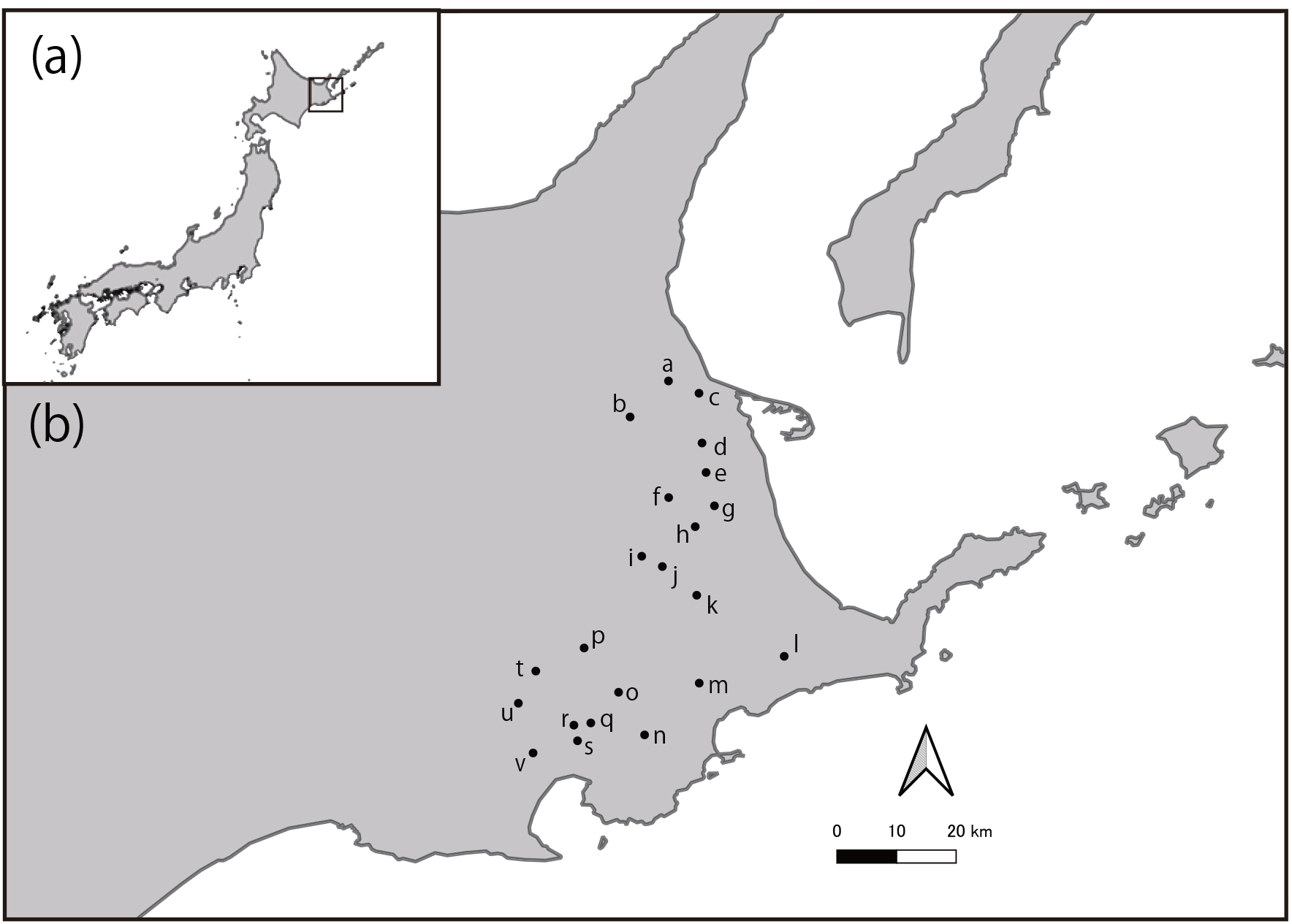
Geographical locations of the study region (a) and representative points of 22 rivers (sites) (b). The actual locations of the study reach on the streamlines and names of the studied rivers remain undisclosed for target species conservation.

### Recruitment status assessment

Quadrat-based quantitative and semi-quantitative mussel sampling was conducted from June to September 2017. Two study reaches (mean ± SD, 27.40 ± 6.17 m), which were apart from each other >500m, were established at each site. Five transects were laid at equal intervals in each reach and three 0.09-m^2^ quadrats were established on each transect. On a single transect, each quadrat was established separately on both sides near the bank and at the mid-channel. Mussels were thoroughly collected from each quadrat using a clear-bottomed viewing bucket and a depth of ∼10-cm-bed-sediment excavation with sieving all materials through a 2-mm mesh sieve. At least 100 *Margaritifera* individuals were collected from each study site. When the number of *Margaritifera* individuals collected by quadrat sampling did not reach 100 at the study site, additional mussel collections were conducted until a total of 100 or more individuals were obtained. One investigator then randomly walked and found mussel individuals within the study reach by using a clear-bottomed viewing bucket. A single quadrat was established at the found individuals as the center, and mussels were collected using a previously described method. The collected mussels were photographed in the field for size measurement and were released to the original study reach. The shell length and height of each mussel were estimated using the ImageJ software (Schneider et al. 2012). The collected mussels were identified using the non-lethal criteria of Miura et al. (2019b). A histogram of the size structure of each site was drawn, and then the data from the two study reaches were combined.

The proportion of ≤ 10-year-old individuals within the population (fraction, “the proportion of juveniles” hereafter) was calculated and used as an index of recruitment success (Österling et al. 2008). The threshold sizes of 10-year-old individuals were estimated based on the growth model following the protocol used by Kawajiri et al. (2021) (see Supplinfo 1).

### Host-fish-availability assessment

To determine host density, 2-pass electrofishing was carried out at each site using a performed using a backpack electrofisher (Model 12 B, Smith-Root Inc., Vancouver, WA, USA) from August to early September 2017 and 2018. The sampling season was consistent with the parasitism period of *M. laevis* larvae in the study region (Kurihara and Goto 2011). At each site, a single reach was established near the mussel sampling location, with an approximate width that was 10 times greater than the mean wetted width. The mean wetted width at each site was calculated by averaging the wetted width values measured at five transects placed at equal intervals within the reach of the fish collection. The total number of host fish collected was counted at each site. All collected host fish were measured for fork length (FL, mm) after anesthetization (FA100, DS Pharma Animal Health Co., Osaka, Japan). Young-of-the-year (YOY) individuals are often crucial hosts for freshwater mussel recruitment owing to their lack of acquired immunity (Österling et al. 2008). Thus, the host density (N 100 m^-2^) was calculated using only the data from the YOY individuals. YOY and other age classes were distinguished based on FL (Fig. S1 in Supplinfo 2). However, two collected *O. masou masou* individuals escaped just before the size measurements. These two individuals were included in the data to calculate host density.

### Statistical analyses

Generalized linear models (GLMs) with binomial error distribution were constructed to test the relationship between recruitment performance and host-fish availability. The proportion of juveniles was used as the response variable and host density was used as an explanatory variable for model construction. Model significance was tested using the likelihood ratio test against the null model (i.e., with no explanatory variables). All statistical analyses were performed by R 4.2.2 (R Core Team 2022) with a significant level of *a* = 0.05. The models were constructed using glmmTMB v.1.1.5 (Brooks et al. 2017).

## Results

A total of 2,623 *M. laevis* individuals (range:24–352 [mean ± SD:119.23 ± 93.77]) were collected. Thirteen sites (59.1%) showed no signs of recent recruitment within 10 years (Fig. 2). The proportion of juveniles ranged from 0.00–0.09 (0.02 ± 0.03) fraction. The size structures of the studied rivers showed that most populations had a predominance of more than 50 mm shell length, with a peak of 50–100 mm size (Fig. 2). The *M. laevis* density was 1.11–130.00 (28.33 ± 40.98) individuals m^-2^, which had no clear relationship with the proportion of juveniles (*r* = 0.04, *P* = 0.86).

**Fig. 2.**
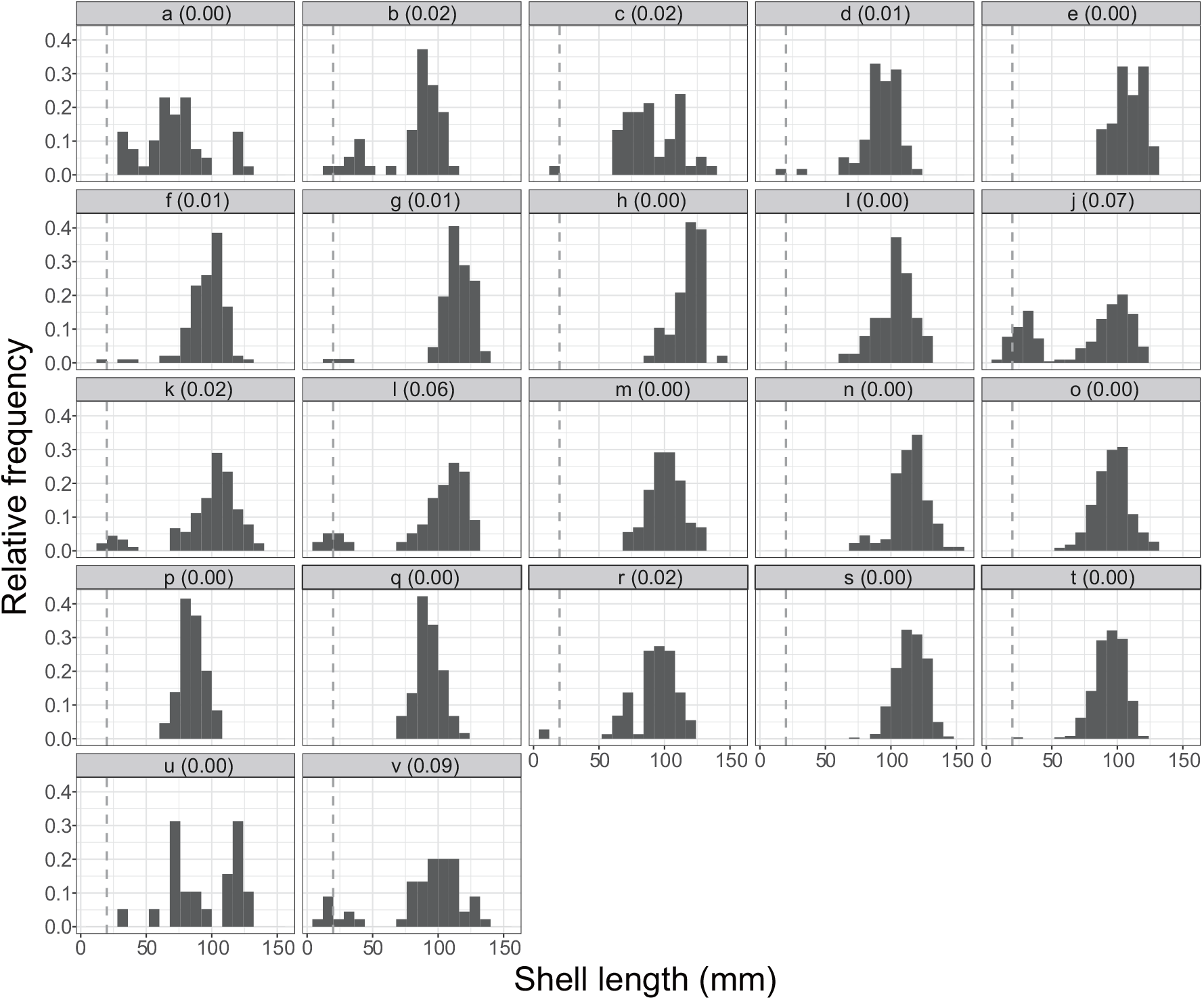
Relative frequencies of shell length of *Margaritifera laevis* in the 22 study rivers. Site names and the proportion of ≤ 10-year-old individuals within the population are shown at the top of each panel. Gray dashed lines indicate the shell sizes of 10-year-old individuals.

A total of 437 YOY host fish (0–113 [19.86 ± 31.75]) were collected by electrofishing. No host fish were collected from five of the 22 sites (rivers *k, n, q, u*, and *v*). The host density ranged from 0.00 to 13.04 (3.50 ± 3.92) 100 m^-2^. Host density was positively associated with the proportion of juveniles (Fig. 3). The GLM, including host density as an explanatory variable, was statistically different and more plausible than the null model (AIC:91.42 vs 121.35, *P* < 0.001), indicating that host density was an influential factor for the proportion of juveniles.

**Fig. 3.**
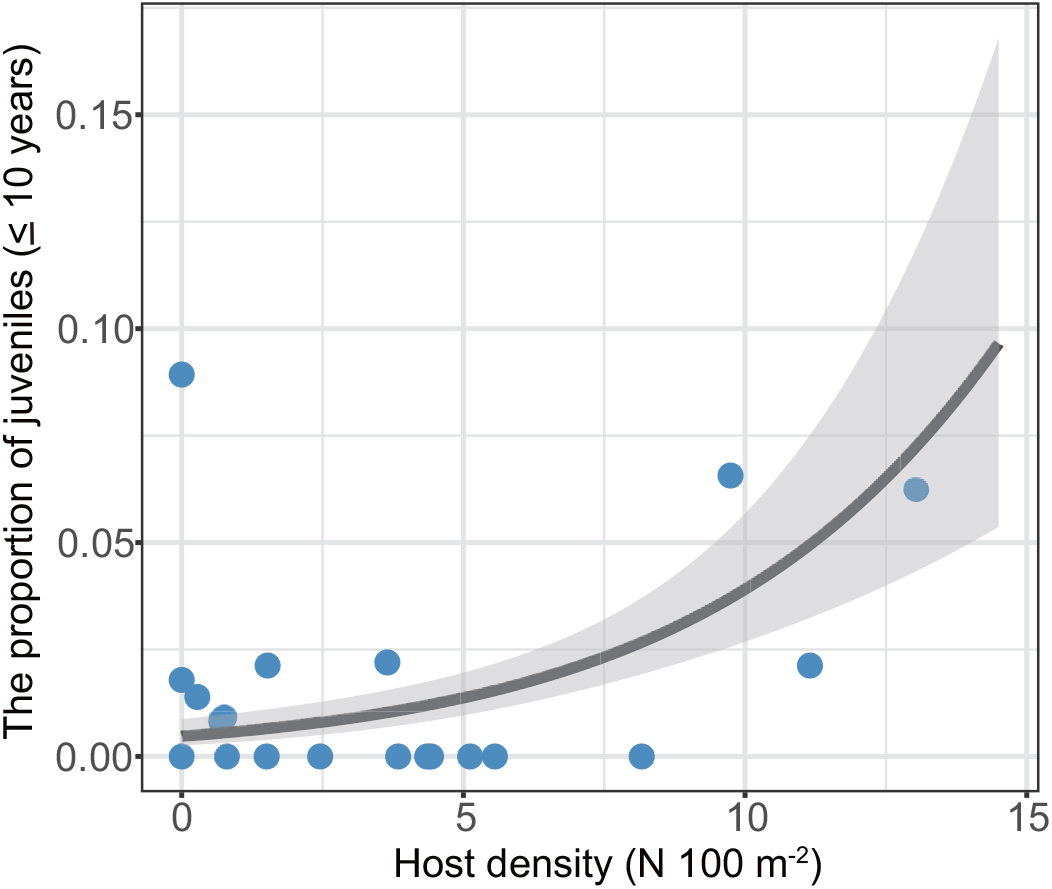
Relationship between the proportion of 10-year-old individuals within the *Margaritifera laevis* population and host density (N 100 m^-2^). The solid curve was drawn by regression of the generalized linear model. Gray-shaded areas indicate 95% CI.

## Discussion

Although *M. laevis* has been examined as an “endangered species” in Japan (Ministry of the Environment of Japan 2020), the recruitment status of this species, together with the evaluation of host-fish availability, have been assessed in very few studies (Akiyama 2007; Kawajiri et al. 2021). Such information on *M. laevis* populations is lacking, particularly in eastern Hokkaido, where a relatively large number of populations have remained (Miura et al. 2019b). Our study is the first to report the recruitment status of *M. laevis* in the study region, and revealed that 59.1% of the population (13 rivers) had no signs of recent recruitment within 10 years among the 22 study rivers. This suggests that these non-recruiting populations are threatened by sustained recruitment failure, which is similar to the recent global trend observed in other freshwater mussel populations (Österling et al. 2008; Geist 2010; Strayer and Malcom 2012). In addition, the present results suggest that host-fish limitation is among the important bottlenecking factors for *M. laevis* recruitment in this region.

Together with existing limited information on *M. laevis* population demographics, our findings improved the current view of *M. laevis* recruitment status. For example, Kawajiri et al. (2021) examined the proportion of 10-year-old juveniles within *M. laevis* populations from 10 rivers in the central, northern, and eastern parts of Hokkaido. Except for one river in eastern Hokkaido (river *j* of the present study), only one river (11.1%) of the nine rivers had no sign of *M. laevis* recruitment with a mean fraction of 0.15 (range:0.00–0.48) of the juvenile proportion. In addition, Akiyama (2007) showed that all 12 targeted rivers in Honshu and the central and northern parts of Hokkaido, Japan, contained ≤ 10-year-old individuals within the *M. laevis* populations. When we compared the present results with those of both studies, the predominance of non-recruiting populations (59.1% of the studied populations) with a low mean proportion of juveniles (0.02 [0.00–0.09] fraction) suggests that *M. laevis* populations in the study region are more seriously threatened by sustained recruitment failure. Thus, greater attention should be paid to future declines in *M. laevis* populations in this region, regardless of the number of remnant populations and each population size.

Several studies have attempted to recover the recruitment of freshwater mussels. However, recovering the natural recruitment of freshwater mussels is still difficult and costly (e.g. Galbraith et al. 2018). Therefore, sustaining recruiting populations would be more feasible than recovering the natural recruitment of non-recruiting populations to effectively sustain freshwater mussel populations. For this reason, higher conservation prioritization should be placed on the nine recruiting populations among the targeted 22 populations. Sustaining and/or recovering the recruitment performance of these populations should be first considered. Subsequently, attempts to recover the natural recruitment of 13 non-recruiting populations need to be considered.

A chain of host-parasite co-decline might be responsible for the recruitment failure of *M. laevis* in eastern Hokkaido. Ishiyama et al. (2021) suggested the density of host fish *O. masou masou* declined due to past farmland expansion in the study region, which supports our finding. Previous studies targeting the same or congeneric mussel species in other regions have also suggested that host density is a key determinant of recruitment success (Arvidsson et al. 2012; Stoeckl et al. 2015). For example, Kawajiri et al. (2021) explored the influential biotic and abiotic factors that explain the recruitment of *M. laevis* in 10 rivers in Hokkaido (including river *j* in the present study). They showed that host density was the most influential factor and suggested that low host density limits the recruitment of *M. laevis*. This study provides evidence that the habitats of the host fish should be managed together when planning to sustain or recover the natural recruitment of *M. laevis* in this region.

The present results suggest that increasing the host density could be effective in recovering *M. laevis* recruitment. In the study region, the density of the host-fish species was mainly impaired by nutrient runoff from farmlands in the catchment, which was interactively affected by the underlying geology (Ishiyama et al. 2021). Therefore, catchment-scale management of nutrient runoff, such as the restoration of wetlands and vegetated buffer strips, could help recover the natural recruitment of *M. laevis* by increasing host-fish availability (Weller et al. 2011; Hansen et al. 2021). In addition, the installation of large wood structures would be effective in increasing host density, because large wood structures can increase in-stream habitat heterogeneity (Nagayama et al. 2009). Furthermore, the (re-)introduction of host fish can increase host density (Galbraith et al. 2018). In particular, it may be more effective during the reproductive season of *M. laevis*.

Overall, our findings suggest an urgent need for conservation measures, including host fish’s habitat management, to sustain *M. laevis* populations in eastern Hokkaido. This report will contribute to the planning of conservation measures for the *M. laevis* populations in this region. In the next step, other limiting factors of *M. laevis* recruitment and their relative importance should be determined in detail to recover *M. laevis* recruitment more effectively. In particular, the post-parasitic juvenile stage could also be a bottleneck for freshwater mussel recruitment together with the parasitic stage (Miura et al. 2021a). Evaluating the performance of this stage and its bottleneck factors is especially important for improving *M. laevis* recruitment. In addition, long-term monitoring programs for both *M. laevis* and host-fish populations are needed to better predict future changes in *M. laevis* population dynamics.

## Acknowledgements

We are grateful to K. Ooue, T. Hasegawa, T. Yamada, M. Kitazawa, D. Nishio, R. Kawanishi and members of the Watershed Conservation and Management Laboratory, Hokkaido University, helped with the field and laboratory work. We also thank the members of the Akkeshi Marine Station, Hokkaido University, for helping with field logistics. This work was partly supported by the Takara Harmonist Fund, Pro Natura Fund, Grant-in-Aid for Science Research of Lake Akkeshi and Bekanbeushi Wetland, JSPS Research Fellow Grant (grant number JP18J12458), and the Environment Research and Technology Development Fund (S15 Predicting and Assessing Natural Capital and Ecosystem Services [PANCES]) of the Environmental Restoration and Conservation Agency, Japan.

## Author contribution statement

Conceptualization: DI, FN, HI, JNN, KM, KK, NI; Data curation: KM; Formal Analysis: KM; Funding acquisition: FN, JNN, KM, KK, NI; Investigation: DI, FN, HI, JNN, KM, KK, NI; Methodology: JNN, KM, KK, NI; Project administration: FN, JNN; Resources: FN, JNN, KM, NI; Software: KM; Supervision: FN, JNN; Validation: KM; Visualization: KM; Writing - original draft preparation: KM; Writing - review and editing: DI, FN, HI, JNN, KM, KK, NI

## Data Availability Statement

Data are available from the corresponding author upon reasonable request.

