## Supplemental information1 for "Recruitment status and host-fish-limitation threats to endangered freshwater pearl mussel (*Margaritifera laevis*) in eastern Hokkaido, northern Japan"


**Author names and affiliations**

Kazuki Miura[1*, 2] ・Nobuo Ishiyama[3, 4] ・Junjiro N. Negishi[5] ・Keita Kawajiri[6] ・Hokuto Izumi[1] ・

Daisetsu Ito[1] ・Futoshi Nakamura[3]

[1] Graduate School of Environmental Science, Hokkaido University, N10W5, Sapporo, Hokkaido 060-0810, Japan

[2] (Present address) Research Institute of Energy, Environment and Geology, Hokkaido Research Organization, N19W12, Sapporo, Hokkaido 060-0819, Japan

[3] Research Faculty of Agriculture, Hokkaido University, N9W9, Sapporo, Hokkaido 060-8589, Japan

[4] (Present address) Forest Research Institute, Hokkaido Research Organization, Bibai, Hokkaido 079-0198, Japan

[5] Faculty of Environmental Earth Science, Hokkaido University, N10W5, Sapporo, Hokkaido 060-0810, Japan

[6] Graduate School of Agriculture, Hokkaido University, N9W9, Sapporo, Hokkaido 060-8589, Japan

***Corresponding author**

Kazuki Miura: Research Institute of Energy, Environment and Geology, Hokkaido Research Organization, N19W12, Sapporo, Hokkaido 060-0819, Japan. Tel.: +81-11-747-3521,

**Age estimation and the results of growth model fittings for *Margatifera laevis***

To determine the threshold size of 10-year-old individuals, growth models were developed following the protocol described in a previous study (Kawajiri et al. 2021). In total, 58 shells of *Margatifera laevis* were collected from the study watersheds between 2017 and 2018. Shell length and height of each individual were measured and identified using the criteria described by Miura et al. (2019). For each individual, the number of growth rings on the ligament was counted under a stereomicroscope. The growth rings on the abraded section of the ligament were estimated by the function of ligament length and the growth ring numbers obtained from 13 non-eroded shells of *M. laevis* juveniles (shell length:9.7–22.8 mm), as described by Kawajiri et al. (2021). The growth curves of *M. laevis* were obtained by fitting four non-linear models: Bertalanffy growth function, Hyperbolic saturation function, Logistic function, and Gompertz function. We then used the measured shell length and counted the growth ring numbers (i.e., age) of *M. laevis* individuals. (San Miguel et al. 2004; Akiyama and Iwakuma 2010; Haag and Rypel 2011; Nakamura et al. 2018). The four models are defined as follows.

Bertalanffy growth function

$$L_t = L_\infty(1 - e^{-k(t-t_0)})$$

Hyperbolic saturation function

$$L_t = \frac{L_\infty k(t - t_0)}{1 + k(t - t_0)}$$

Logistic function

$$L_t = \frac{L_\infty}{1 + e^{-k(t-b)}}$$

Gompertz function

$$L_t = L_\infty e^{-ae^{-kt}}$$

where $L_\infty$ is the theoretical maximum length (or asymptotic length, mm), $k$ is the growth coefficient (year$^{-1}$), $t_0$ is the theoretical age at zero length (years), and $a$ and $b$ (years) are constants (Akiyama and Iwakuma 2010). The best-fitting model was selected by comparing RSS (the residual sum of squares (RSS) (Akiyama and Iwakuma 2010). All growth model construction was performed using R 4.2.2 (R Core Team 2022) with a package of nlsMicrobio v. 0.0–3 (Baty and Delignette-Muller 2013).

As a result of the comparisons of the four growth model fittings, Hyperbolic saturation function was the best-fitted function with the lowest RSS (Table A and Fig. A). The estimated shell length of 10-year-old individuals was 19.89 mm.

Table A. Parameter estimates of four growth functions for *M. laevis* in eastern Hokkaido.

| **Function** | **Parameters** | | | | | |
|---|---|---|---|---|---|---|
| | $L_\infty$ | $k$ (year$^{-1}$) | $a$ | $b$ | $t_0$ | **RSS** |
| **Bertalanffy** | 120.52 | 0.04 | - | - | 4.91 | 3499 |
| **Hyperbolic saturation** | 160.41 | 0.03 | - | - | 5.75 | 3292 |
| **Logistic** | 102.75 | 0.11 | - | 21.66 | - | 5269 |
| **Gompertz** | 108.44 | 0.07 | 3.28 | - | - | 4310 |

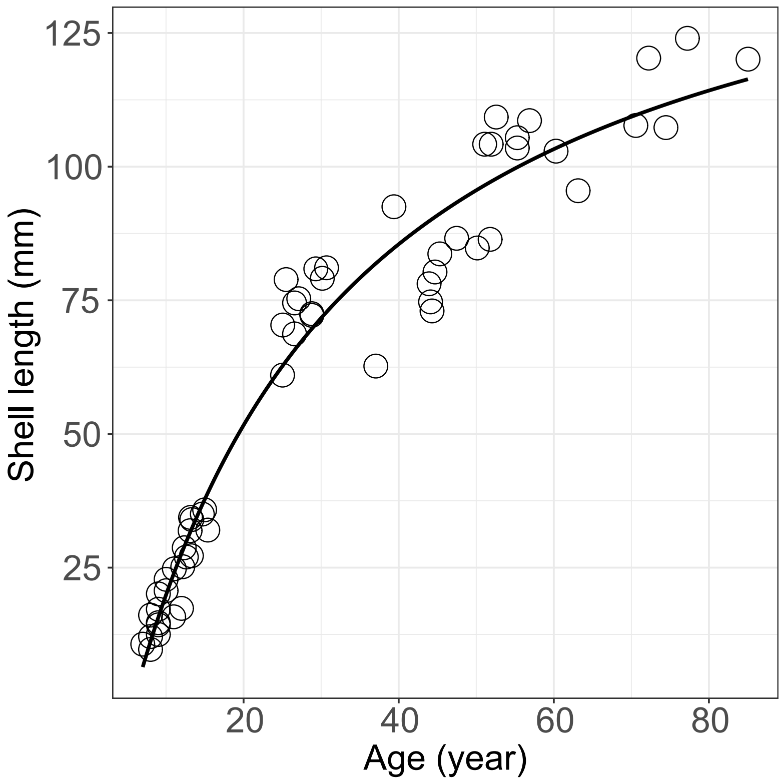


Fig. A Biplot showing the relationship between the age and shell length of *M. laevis* and the estimated Hyperbolic saturation growth curve.
