## Supplemental information2 for "Recruitment status and host-fish-limitation threats to endangered freshwater pearl mussel (*Margaritifera laevis*) in eastern Hokkaido, northern Japan"


**Author names and affiliations**

Kazuki Miura[1*, 2] ・Nobuo Ishiyama[3, 4] ・Junjiro N. Negishi[5] ・Keita Kawajiri[6] ・Hokuto Izumi[1] ・

Daisetsu Ito[1] ・Futoshi Nakamura[3]

[1] Graduate School of Environmental Science, Hokkaido University, N10W5, Sapporo, Hokkaido 060-0810, Japan

[2] (Present address) Research Institute of Energy, Environment and Geology, Hokkaido Research Organization, N19W12, Sapporo, Hokkaido 060-0819, Japan

[3] Research Faculty of Agriculture, Hokkaido University, N9W9, Sapporo, Hokkaido 060-8589, Japan

[4] (Present address) Forest Research Institute, Hokkaido Research Organization, Bibai, Hokkaido 079-0198, Japan

[5] Faculty of Environmental Earth Science, Hokkaido University, N10W5, Sapporo, Hokkaido 060-0810, Japan

[6] Graduate School of Agriculture, Hokkaido University, N9W9, Sapporo, Hokkaido 060-8589, Japan

***Corresponding author**

Kazuki Miura: Research Institute of Energy, Environment and Geology, Hokkaido Research Organization, N19W12, Sapporo, Hokkaido 060-0819, Japan. Tel.: +81-11-747-3521,

**Supplemental figure**

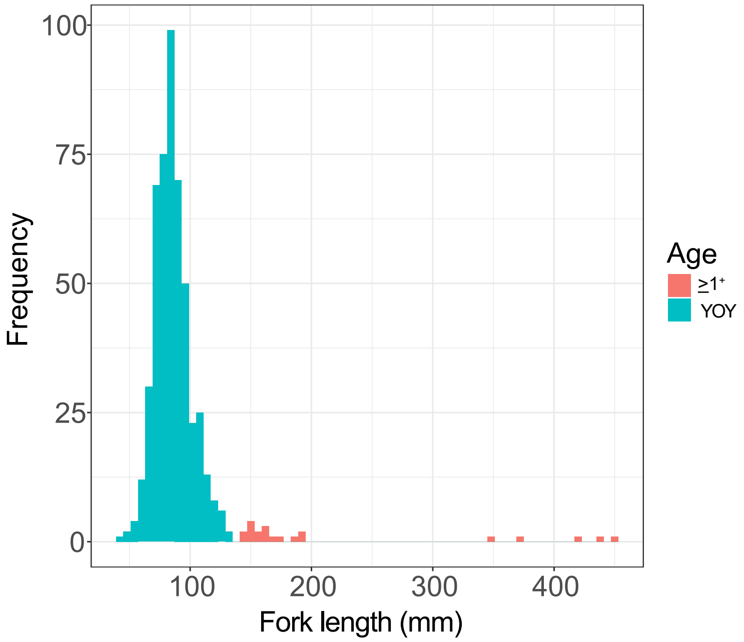


Fig. S1 Fork length (mm) distribution of collected host fish (*Oncorhynchus masou masou*). Blue and red indicate young-of-the-year (YOY) and $\geq 1^{+}$ individuals, respectively.
